# Apolipoprotein interaction induces shape remodeling and lipid phase separation in giant unilamellar vesicles

**DOI:** 10.1101/2025.04.10.648283

**Authors:** Christopher F. Carnahan, Wei He, Tugba N. Ozturk, Yaqing Wang, Viviane N. Ngassam, Aleksandr Noy, Timothy S. Carpenter, John C. Voss, Matthew A. Coleman, Atul N. Parikh

**Affiliations:** Biophysics Graduate Group, University of California, Davis, CA; Biosciences and Biotechnology Division, Lawrence Livermore National Laboratory, Livermore, CA; Materials Science Division, Lawrence Livermore National Laboratory, Livermore, CA; Department of Biomedical Engineering, University of California, Davis, CA; Department of Biochemistry & Molecular Medicine, University of California, Davis, CA; Institute for Digital Molecular Analytics & Science, Nanyang Technological University, Singapore

**Keywords:** Self-assembly of lipid nanodiscs, lipoproteins, protein0lipid interactions, giant unilamellar vesicles

## Abstract

Apolipoprotein A-I (ApoA-I) – a 243-residue amphipathic protein containing an N-terminal globular domain and a primarily helical C-terminal lipid binding domain – is a principal protein component of high-density lipoprotein (HDL) or “good” cholesterol, which is an essential component of lipid homeostasis in humans. Synthesized in the liver and intestine and excreted in the blood, ApoA-I undergoes complex, cooperative, and dynamic self-assembly with membrane lipids, producing unlipidated (or weakly lipidated), nascent discoidal, and mature HDL states. In vitro studies demonstrate that the reconstitution of purified protein and lipids restores this cooperative self-assembly. However, the kinetic pathways by which these mesoscopic, proteolipidic assemblies form remain incompletely understood. Here, we monitor the dynamics of ApoA-I-membrane interactions through real-time monitoring of morphological changes, which ensue when ApoA-I is incubated with minimal giant unilamellar vesicles (GUVs) composed of single phospholipids or phase-separating phospholipid-cholesterol mixtures. Our fluorescence microscopy measurements reveal that the interaction initiates a gross, morphological remodeling of the parent vesicle proceeding through discrete stages involving membrane poration, solute leakage, vesiculation, and lipid-lipid phase separation. Our atomic force microscopy measurements confirm that the outcome includes discoidal nanoparticles. This qualitative phenomenology is robust and fully reproducible for different protein mutants and alleles (WT APOA-1, Δ49ApoA-I, ApoE-3, and ApoE-4) and other lipid mixtures (including mixtures containing phosphoserine lipids). Our molecular simulations recapitulate the essential shape changes and further reveal the composition dependence of the interactions. Together, these findings outline key steps in protein-lipid interactions that facilitate the assembly of mesoscopic reconstituted lipoproteins and nanodiscs.

## INTRODUCTION

Lipids are amphiphilic molecules that play essential roles in membrane synthesis, energy storage, and cell signaling. The ability to transport them from the sites of absorption (or syntheses) to other cells and tissues – as a means to distribute metabolic energy – represents a critical step in the evolution of multicellular organisms (*1*). In animals, this function is enabled by the collective transport of lipids in the form of micelle-like, colloidal nanoparticles through the aqueous circulatory system.

Collectively termed lipoproteins, these circulating nano-particles are supramolecular, mesoscopic complexes consisting of mixtures of several different classes of lipids (e.g., triglycerides, cholesterol, and phospholipids) and one or more proteins called apolipoproteins (*2*). Morphologically, they are organized into spheroidal particles consisting of an outer layer of phospholipids, unesterified cholesterol, and proteins, encapsulating the central core made up of neutral lipids, predominately cholesteryl ester, and triacylglycerols (*3*). Although most lipoproteins share these structural features, they also exhibit considerable structural, compositional, and morphological variations. An early basis for the classification and fractionation of the lipoprotein diversity was the particle density: different lipoproteins floated differentially in repeated ultracentrifugations with progressively increased densities (*4*). Sub-sequent efforts revealed additional criteria for classifying lipoprotein diversity (*5, 6*), including sizes, electrophoretic mobilities, lipid and protein compositions, and metabolic functions. Currently, the most common classification recognizes six different classes of human lipoproteins (*2*). These classes include (i) *chylomicrons*, which are >100 nm in diameter and commonly synthesized in intestine; (ii) *chylomicron remnants*, which form after lipoprotein lipase hydrolyze the triglycerides of the source chylomicrons; (iii) *very-low-density lipoproteins* (VLDL, d < 1.006 g/mL and diameter, 30-90 nm), produced in the liver; (iv, v) *intermediate-density lipoproteins* (IDL, d = 1.006-1.019 g/ml) and *low-density lipoproteins* (LDL, d = 1.019-1.063 g/mL).), produced by the lipase activity on VLDLs; and (6) *high-density lipoproteins* (HDL, d = 1.063-1.21 g/mL), which have the smallest sizes of 8-12 nm in diameter. Within each of these classes, a significant compositional variability exists, making lipoproteins one of the most compositionally heterogeneous classes of particles in blood.

In addition to being the smallest (∼8-12 nm in diameter) and the densest (d = 1.063-1.21 g/mL), HDLs are also compositionally and functionally distinct (*7*). Metabolically, they play a major role in reverse cholesterol transport, a mechanism by which circulating HDLs draw cholesterol from the cells of the peripheral tissues and return it to the liver for excretion (*8, 9*). This role in mediating macrophage cholesterol efflux is thought to be primarily responsible for the inverse correlation between their concentrations in plasma levels and the risk for coronary artery disease (*10, 11*), rendering HDLs uniquely atheroprotective. HDLs are also compositionally distinct from their lipoprotein counterparts. Their lipids are not solubilized by apolipoprotein B. Instead, they are bounded, most dominantly, by ApoA-I – a single polypeptide of 243 amino acids secreted in the intestine and liver (*12, 13*) – which accounts for ∼70% of the total protein mass of HDL particles (*7*).

*In vivo*, HDL assembly begins with the activity of ATP-binding cassette transporter ABCA1, a membrane protein. According to the alternating access mechanism, an activated ABCA1 flips phospholipids from the cytoplasmic to the outer, exocytoplasmic leaflets of plasma membranes (*14*). The resulting excess lipid density (and area) in the outer leaflet promotes vesiculation and efflux of phospholipids and cholesterol to the extracellular space in the form of vesicles. A recent computer simulation (*15*) proposes an alternate view. It suggests that upon activation, ABCA1 acts as a phospholipid translocase, extracting lipids from the outer exocytoplasmic leaflet to form a luminal structure, that migrates into the extracellular space as vesicles. In both cases, lipid-poor ApoA-I activates ABCA1 and interacts with the vesicles produced by the ABCA1 activity in the extracellular space, leading to the formation of nascent HDLs. These newly formed HDLs are discoidal in morphology. Subsequent esterification of cholesterol by the enzyme lecithin: cholesterol acyltransferase (LCAT) initiates the formation of the hydrophobic core and induces the shape transformation from the discoidal to the spheroidal morphology (*16*).

As discrete nanoscale compartments transporting lipids and membrane proteins through aqueous environments, HDLs and their synthetic counterparts are appealing for many biomolecular technologies (*17*). Reconstituted *de novo* by incubating pre-determined mixtures of lipids with apolipoprotein (as well as their truncated variants called membrane scaffold proteins or MSPs) or synthetic polymers (e.g., styrene-maleic acid, SMA) (*18, 19*), these reconstituted HDLs (rHDLs) – also called nanolipoproteins (NLPs) and nanodiscs – mimic an intermediate discoidal state of ApoA-I found *in vivo* (*20*). These constructs are proving valuable for solubilizing membrane proteins (*21- 24*), tailoring membrane compositions (*25*), and screening membrane-protein targeting drugs and therapies (*26, 27*).

For both the biotechnological and biomedical applications of HDLs, understanding molecular mechanisms by which ApoA-I disrupts topologically closed lipid vesicles is important. Much of our current knowledge is obtained from computer simulations and *in vitro* reconstitution studies. Two limiting modes of protein-lipid association in the formation of discoidal rHDLs have been proposed: (i) the picket fence model, where two ApoA-I molecules wrap around a disc of a single lipid bilayer with adjacent alpha-helices in antiparallel orientation, parallel to the acyl chain of the phospholipids (*28, 29*) and (ii) the double-belt model, where two antiparallel ApoA-I are oriented perpendicular to the lipid tails (*30*). The bulk of the experimental evidence currently favors the belt model; however, the hybrid conformations in HDL subpopulations, where some of the ApoA-I helices are perpendicular to the lipid tails, have yet to be ruled out.

For either model, the dynamics of the alpha-helical segments of ApoA-I play important roles. ApoA-I is an amphipathic protein that contains a globular N-terminal domain (residues 1-43) and a lipid-binding C-terminal domain (residues 44-243). As secreted, lipid-poor ApoA-I features five amphipathic helices spanning residues 7-44, 54-65, 70-78, 81-115, and 147-178. Together, these helices make up approximately 50% of the protein content (*31*). The free energies of stability for these helices in the lipid-free state are estimated to be in the range of 3-5 kcal/mol (∼5-9 *k*_*b*_ *T*), sufficient to provide stable helical segments, however, these values are lower than 5-10 kcal/mol (∼9-17 *k*_*b*_ *T*) typically observed for globular proteins, suggesting that the protein is highly conformationally dynamic. Together, these features allow ApoA-I to unfold and refold readily. This structural plasticity is essential to its function, allowing for the stability of both the discoidal and spheroidal assemblies with varied lipid stoichiometries. Specific protein conformational changes – specifically the increase in the alpha-helical content of the protein (*32, 33*) – that accompany HDL assembly are now well-understood (*34*).

Still, our understanding of how ApoA-I interacts with organized lipid mesophases, such as membranes, vesicles, and liposomes, remains incomplete. Computer simulations and *in vitro* reconstitution studies suggest roles of packing density (*35*), packing defects (*36*), and local membrane curvatures in driving the assembly (*37*). However, the kinetic pathways and structural intermediates – how ApoA-I invades a topologically closed membrane, disrupts its structure, assembles lipoproteins, and what remains of the parent membrane – remain incompletely understood. To address these questions, we investigate the real-time interactions between a truncated isoform of ApoA-I and lipid bilayers organized as giant unilamellar vesicles. By employing a platform that enables direct visual characterization, we identify previously unappreciated biophysical steps – including poration, vesiculation, and phase separation – which mark the ApoA-I – membrane interactions *en route* to the formation of HDL-like particles.

## RESULTS

We begin by investigating the interaction between ApoA-I and single lipid bilayers. For our study, we used a N-terminally truncated isoform of ApoA-I in which the first 49 amino acids are removed: Δ49ApoA-I. This modification has been previously shown to preserve the essential lipoprotein-lipid interactions (*38*). This modification to the N-terminal helical bundle may influence the proteins lipid specificity, as this region is believed to be involved in its essential function of lipid binding (*39*). Our lipid source for reconstituting HDL-like particles is a suspension of protein- and cytosol-free, model phospholipid membranes of closed topologies, i.e., giant unilamellar vesicles (GUVs, 15-30 µm diameter) (*40*). We used both single-component and multicomponent GUVs. Our single component GUVs consist of the mono-unsaturated phospholipid, 1-palmitoyl 2-oleoyl-*sn*-1-glycero-3-phosphocholine (POPC). Our multi-component GUVs are reconstituted using ternary lipid mixtures composed of POPC, egg-sphingomyelin (SM), and cholesterol (Chol) in equimolar (1:1:1) and cholesterol poor (2:2:1) proportions. Depending on composition and temperature, these lipid mixtures are known to organize as a singular, uniform phase or exhibit phase separation (*41*), forming two co-existing liquid phases: a dense phase enriched in SM and Chol, designated as the liquid-ordered phase (L_o_ ) and a second, less dense liquid-disordered phase (L_d_ ) consisting primarily of POPC. Differentiation between the L_o_ and L_d_ phases through fluorescence microscopy is achieved by doping the GUVs with small concentrations (1 mol %) of a phase sensitive probe, N-lissamine rhodamine dioleoyl phosphatidylethanolamine (Rho-DOPE, Ex/Em, 560/583 nm) (*42*). At the equimolar (1:1:1) composition, the phase diagram predicts the absence of domain formation at optical length scales (*41*) but does not rule out existence of nanoscale domains (*43*). Consistent with this prediction, GUVs encapsulating 100 mM sucrose appear optically homogeneous at room temperature in an osmotically balanced, isotonic medium containing 100 mM glucose. Monitoring the temporal changes using spinning disc confocal and wide-field fluorescence microscopy allows us to follow the morphological and topological transformations of these GUVs.

Introducing Δ49ApoA-I to a suspension of single component, POPC GUVs (decorated with 1 mol % Rhodamine PE and encapsulating 100μM of an NBD-labeled glucose (NBD-G, Ex/Em, 475/550 nm) in the interior) triggers gross membrane remodeling (**Supplementary Video 1**) characterized by a well-defined sequence of structural, morphological, and topological transformations (**Fig. 1**).

**Figure 1:**
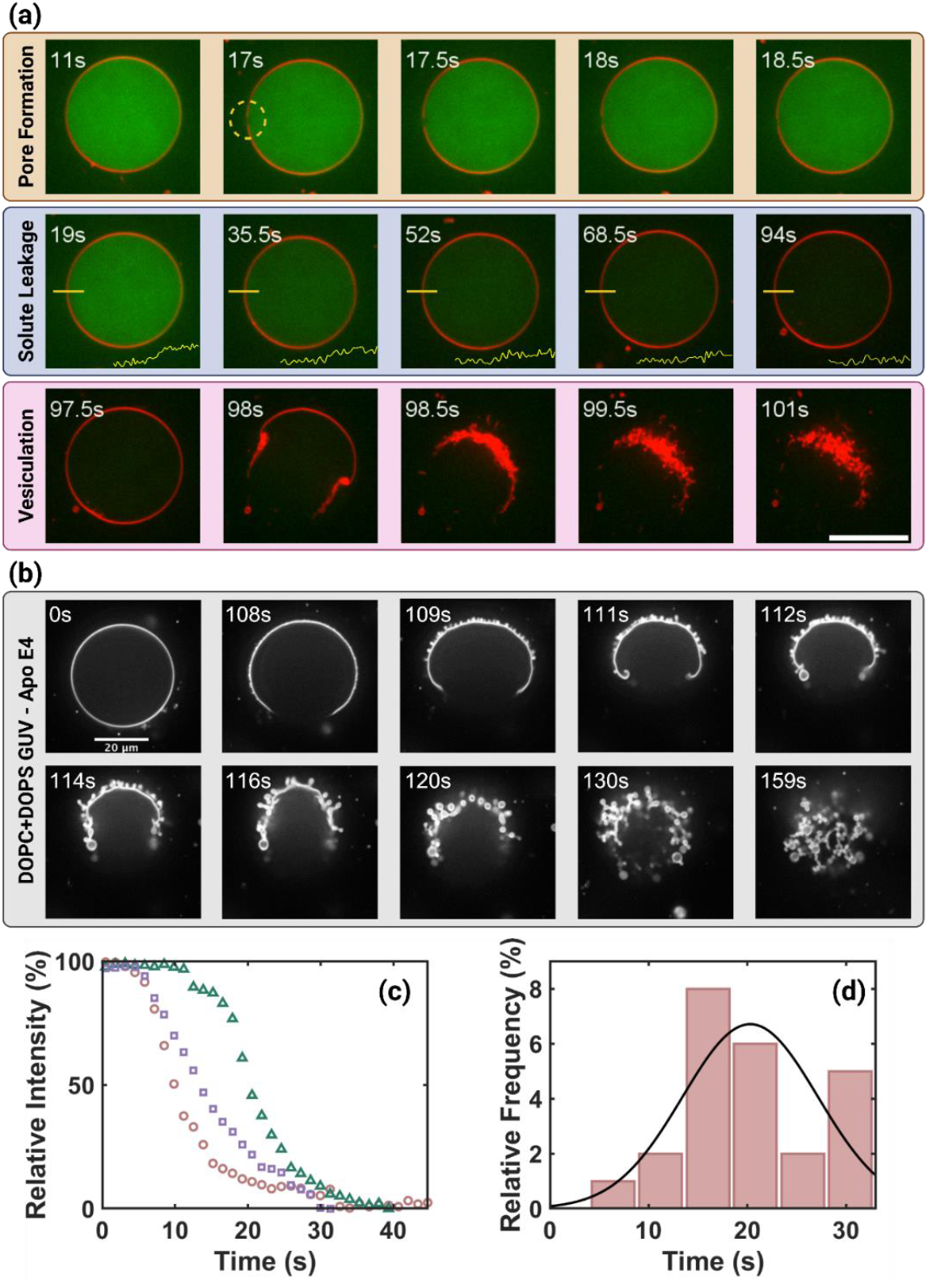
Real-time visualization of membrane-apolipoprotein interactions. (a) A montage of images from a time-lapse video of confocal fluorescence microscopy images of single giant unilamellar vesicles (GUVs) in aqueous suspensions interacting with Δ49ApoA-I. The GUVs consist of POPC doped with 1 mol % Rho-B-DOPE (red) and encapsulate 100µM of a soluble, membrane impermeable dye NBD-G (green). The suspension is exposed to 0.88µM Δ49ApoA-I (Supplementary Video 4). The images are selected to highlight the three major morphological events, namely pore formation, solute leakage, and vesiculation, which characterize the dynamics. The yellow bar is the line along which pixel intensities are measured. Scale bar 10µm. (b) Corresponding montage for GUVs consisting of mixtures of DOPC, DOPS, and Rho-PE (79:20:1) GUV exposed to 20µM ApoE4. (c) Leakage profiles for NBD-labeled glucose from POPC GUVs (n=3)following the addition of ∼0.88 µM Δ49ApoA-I (supplementary video 2) (d) A histogram for full leakage of NBD-G from POPC GUVs (n=24 GUVs) (t^-^ =20.3s).

**Figure 1:**
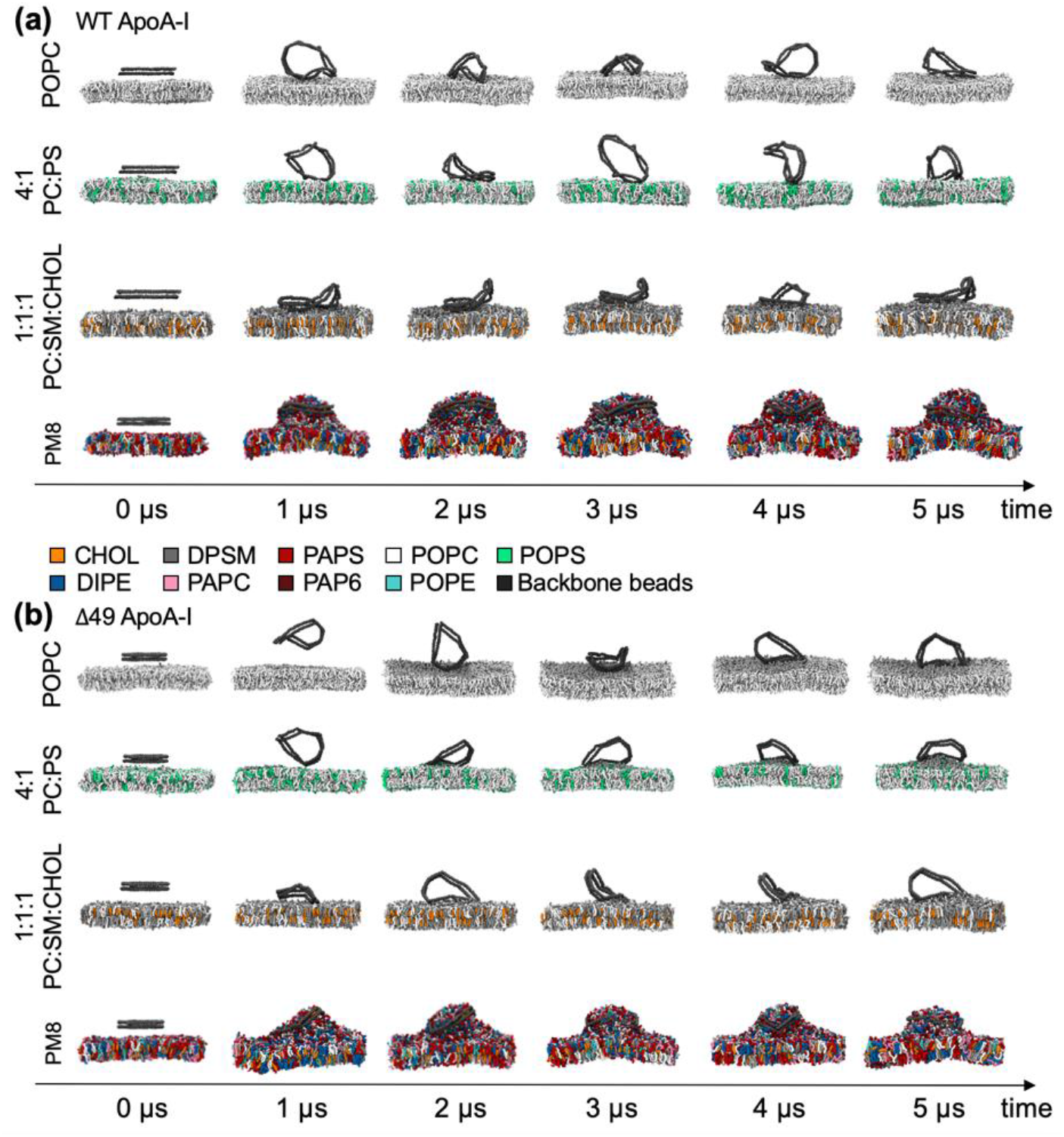
Coarse-grained molecular dynamics simulations of the WT andΔ49 ApoA-I. The snapshots of **(a)** the WT ApoA-I and **(b)Δ**49 ApoA-I double-belt proteins in four different membranes are taken at every 1 µs throughout a single representative 5 µs-long simulation trajectory as indicated by the arrow; the first column of each row corresponds to the initial placement of the pre-assembled double-belt protein 4 nm away from the membrane surface. In the panels a and b, each row corresponds to a system containing a different membrane composition: from top to bottom, the systems include a 100% POPC, 4:1 POPC:POPS, 1:1:1 CHOL:DPSM:POPC and PM8 (mammalian plasma membrane model) membrane. The representative simulations shown here are carried out with the restraints ensuring double-belt proteins remain open and flexible. Lipids are color-coded: CHOL in orange, DIPE in blue, DPSM in gray, PAPC in pink, PAPS in red, PAP6 in dark red, POPC in white, POPE in cyan and POPS in green. Water and ions are not shown for clarity. Protein backbone beads are colored in dark gray.

### Δ49ApoA-I-membrane interactions produce microscopic pores

The first major event involves a transient, short-lived change in intensity at the membrane surface. Here, the uniform fluorescence intensity decorating the equatorial plane in the confocal fluorescence images of single GUVs is transiently replaced by a Janus binary pattern of fluorescence (**Fig. 1a - 17s**). The intensity pattern consists of (i): a dominant segment with the fluorescence intensity comparable to that before the change and (ii) a smaller arc or a domain, roughly 2 µm in diameter, with a vanishingly low fluorescence intensity comparable to the background value (I_membrane_ / I_arc_ = 1,400 %). This intensity perturbation lasts less than 2 seconds before returning to its original state. Comparing across multiple experiments, we find that the event is reproducible; it also exhibits considerable variability in the lag time (10-500 s), time after the introduction of protein that we witness the event, and the lifetime (200 ms – 2 s), the time over which patterned intensity persists (**Figure S1**). In all cases, these short-lived arcs of low fluorescence are of microscopic dimensions (2-5 µm), reminiscent of a transient microscopic pore in the GUV membrane resulting from an osmotic imbalance. The variable lag time we observe is consistent with the stochastic nature of pore formation.

### Solute leakage through a cascade of pore formation events

A key consequence of the formation of transient pores is the mixing of the GUV-encapsulated solution with the external bath. To track this mixing, we monitor the leakage of a membrane-impermeable fluorescent probe, NBD-glucose (green, 100 µM, permeation constant, ∼6-25 x 10_-9_ cm/s) (*44*), pre-encapsulated within the GUVs. Images shown in **figure 1a** reveal that the fluorescence intensity of the dye gradually diminishes, until optical homogeneity is achieved with the surrounding bath. Observing several different GUVs of different sizes (r = 3-23 μm), we find that the intensity changes of single GUVs follow a sigmoidal decay (**Fig. 1c**) with variable lag time (∼10-700 s) and persists over a broad window of time (5-94 s) before reaching a homogeneous plateau (averaging ∼20 ± 7 s (**Fig. 1d**)).

### Vesiculation and lysis of single-component GUVs

The observations of membrane poration, solute, and solvent exchanges above precede the most dramatic structural remodeling leading to large-scale morphological and topological transformations because of the interactions of GUVs with Δ49ApoA-I. Beginning at ∼98 s (**Fig. 1a**), the epifluorescence images at the equatorial plane of this POPC GUV show a dramatic membrane restructuring. The optically uniform and continuous circle of fluorescence at the equatorial perimeter is replaced by an open arc terminating into a string of closely spaced, smaller circular features with diameters of less than one micrometer (< 1 µm). These features form abruptly over 2 - 3 frames (< 1 s). The fluorescence emission from these clusters is more intense (I_small_ /I_mother_ = 200%) than that of the surrounding mother membrane. **Supplementary video 2** depicts an example where a similar topological change occurs in nearly all POPC GUVs during the observed period (17 min). These observations are reproducible (**Supplementary Videos 1-6**) across multiple independent experiments.

To determine whether these observations were unique to the specific ApoA-I mutant we tested, we carried out additional experiments using three different apoproteins: wild-type ApoA-I, ApoE-4, and ApoE-3. When these proteins are incubated with GUVs composed of 4:1 DOPC-DOPS, qualitatively similar results are observed. For example, the morphological changes of a 4:1 DOPC-DOPS GUV exposed to 20 µM ApoE-4, respectively is depicted in **Figure 1b**. Additional independent examples of morphological changes following the incubation of comparable 4:1 DOPC-DOPS GUVs with 0.44 µM ApoE-4 and 7.4 µM ApoE-3 are shown in **Figures S3** and **S4**, respectively. Together, these observations confirm that these instances of membrane restructuring are not limited to the WT ApoA-I isoform or its Δ49 variant. Instead, they reflect a consistent pattern of membrane deformations characterizing the interaction between apolipoproteins and lipid membranes.

### Lipid removal and formation of NLP-like structures by Δ49ApoA-I

It is well established that apolipoproteins readily extract lipids from organized membranes to form apolipoprotein structures (NLPs), or nanodiscs (*38, 45*) To directly test whether the incubation of GUVs with Δ49ApoA-I produces these NLP particles, we investigated the GUV bathing medium (purified with size-exclusion chromatography (SEC)) using high-speed atomic force microscopy (HS-AFM). Scanning the deposits of the solution on planar substrates (see experimental methods), we find unambiguous evidence for the appearance of a population of discrete disk-shaped structures (**Fig. 2a**). These structures have an average height of 4.5 ± 1.1 nm and an average diameter of 14.0 ± 4.0 nm (**Fig. 2b & 2c**) – in excellent agreement with the formation of NLPs (*46, 47*) and do not resemble unlipidated protein aggregates (*45, 48*). Note, however, additional independent characterization of the presence of GUV-derived lipids in the observed nanoparticles is needed before an unambiguous assignment can be made.

**Figure 2:**
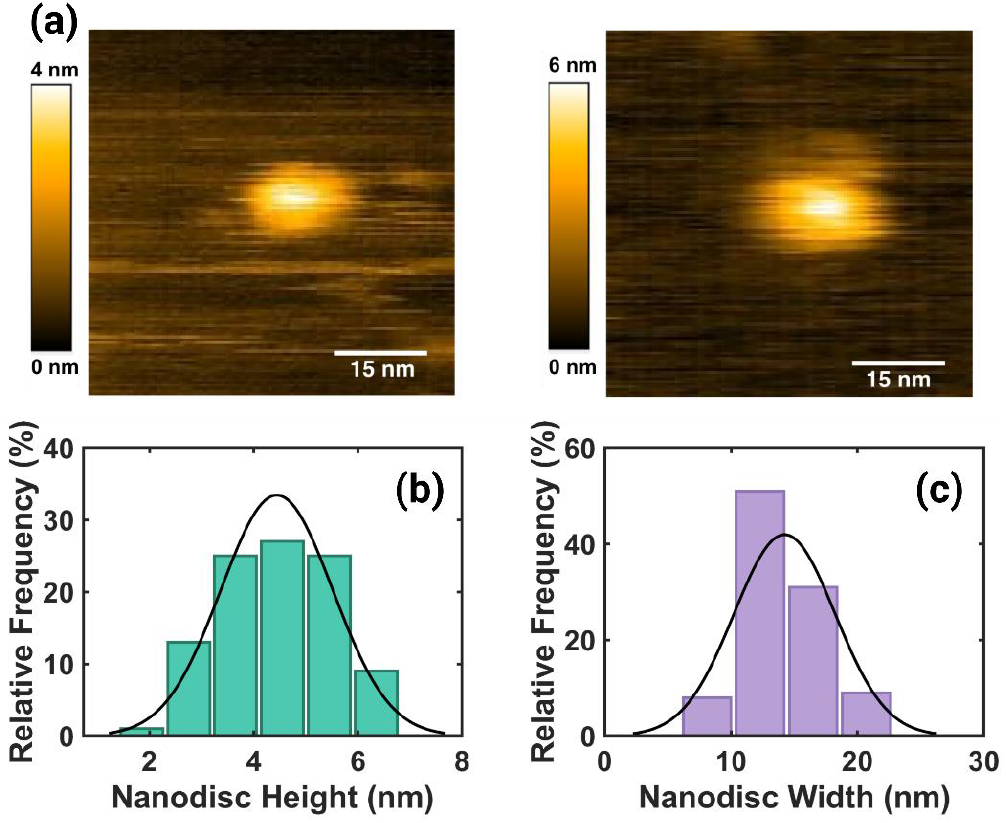
High-Speed atomic force microscopy characterization of nanodiscs. (a) AFM images of representative mesoscopic structures observed by depositing aliquots of the POPC GUV suspension purified by size-exclusion chromatography after incubation with Δ49ApoA-I. Histograms of height and width of typical structures observed.

### Coarse-grained molecular dynamics simulations with WT and Δ49ApoA-I

To further characterize the ApoA-I-mediated membrane deformations, we carried out a series of coarse-grained molecular dynamics simulations. We studied interactions of both the **WT and Δ49ApoA-I** with lipid bilayers composed of 100% POPC, 4:1 POPC:POPS, 1:1:1 POPC:DPSM:Chol and the PM8 lipid mixture (See Methods). For all bilayer compositions, we pre-assembled the double-belt and placed it 4 nm away from the bilayer surface. For each membrane condition, three independent simulations demonstrate that, without any restraints, the double-belt protein collapses into an 8-shape within the first 35-100 ns of the simulations and this conformation remains stable for the rest of the simulations regardless of the membrane composition (**Fig S5. a-d, the first three rows**). With restraints, we ensure that the double-belt conformation remains as an open ring that is still able to undergo significant conformational changes (**Fig S5. a-d, the last three rows**). In 100% POPC membranes (**Fig. S5a**), the WT ApoA-I interacts with the surface, either continuously or intermittently, without significant perturbation in all simulations except for one, in which a small portion of POPC lipid is captured by the protein and remains deformed throughout the rest of the simulation. For mem branes composed of 4:1 POPC:POPS (**Fig. S5b**), at least some parts of the WT ApoA-I continuously interact with the membrane. In 4 of 6 independent simulations containing a 4:1 POPC:POPS membrane, membranes show mild deformations due to interactions with the protein. In 1:1:1 POPC:DPSM:Chol membranes (**Fig. S5c**), the WT ApoA-I intermittently interacts with the membrane, without significant morphological defects. Finally, in the PM8 membranes (**Fig. S5d**), it takes about 300 ns to 1 µs for the WT ApoA-I to fully sit on the PM8 membrane surface (**Fig. 3a**) in all independent simulations and afterwards, stable protein-membrane interactions are maintained. These stable interactions cause significant deformations on the surface of PM8 membranes in all independent simulations. Particularly in the simulations where the protein is restrained to remain flexible and open, the protein causes a bud-like deformation that is the most significant in comparison to the deformations observed in other membrane compositions.

These simulations were repeated with Δ49ApoA-I double-belt (**Fig. 3b**). Similar interactions with the POPC, POPC:POPS, POPC:DPSM:Chol and PM8 membranes were observed. The protein induced the most pronounced bud-like deformations in the PM8 membrane (**Fig. S5d**), and its interactions were subtly disruptive for 4:1 POPC:POPS membranes (**Fig 3b, row 2**) when compared to interactions by WT ApoA-I (**Fig 3a, row 2**).

### Δ49ApoA-I binding-induced phase separation and vesiculation in cholesterol-containing GUVs

The lipid-specific behavior of ApoA-I interactions witnessed during the molecular simulations above raises some general questions: By sequestering and binding specific sub-classes of lipid molecules, does Δ49ApoA-I sort the remaining pool of membrane lipids? Can the protein association break symmetry and transform multicomponent, homogeneous membranes into phase-separated states? These questions are not limited to Δ49ApoA-I but arise in broad classes of protein-membrane interactions where the preferential association of proteins with a sub-set of molecules locally can shift the phase space and drive global phase separation.

To begin to explore the above questions, we incubate ternary GUVs, composed of an equimolar mixture (1:1:1) of POPC, SM, and Chol, with Δ49ApoA-I. To enable visualization of membrane phases by fluorescence, we doped the lipid mixture with small concentrations of two phase-sensitive lipid probes: (i) 1 mol% of Rho-DOPE (red, Ex/Em 560:583), which preferentially associates with the POPC-enriched liquid-disordered phase (L_d_ ), and (ii) 3 mol % NBD-PE (green, Ex/Em 463:536), which selectively partitions within cholesterol- and SM-rich, liquid-ordered (L_o_ ) phase. Previous work has established that the presence of these probe lipids at these concentrations does not influence the phase behaviour of the dominant ternary lipid mixture (*42*).

A selection of images from the time-lapse movies (**Supplementary Video 4 & 5**) shown in **Fig. 4a** and **Fig. 4b** document how Δ49ApoA-I interaction transforms these multi-component GUVs. The GUV is imaged from a distal plane, capturing a confocal view of the apical hemisphere (away from the substrate), which initially appears homogeneous (**Fig. 4a 5:27**). After a short delay, following the addition of Δ49ApoA-1 (8:54), the uniform intensity along the perimeter of the membrane is abandoned, replaced by a pattern of small domains of high intensity of Rho-DOPE (red). In the subsequent several minutes (12:21), membrane domains elevated in NBD-PE intensity (green) form simultaneously with the complementary domains enriched in Rho-DOPE. These segments of the membrane continue to grow and increase in intensity while traversing to and aggregating within the distal hemisphere of the vesicle, characterizing the incipient phase separation.

**Figure 4:**
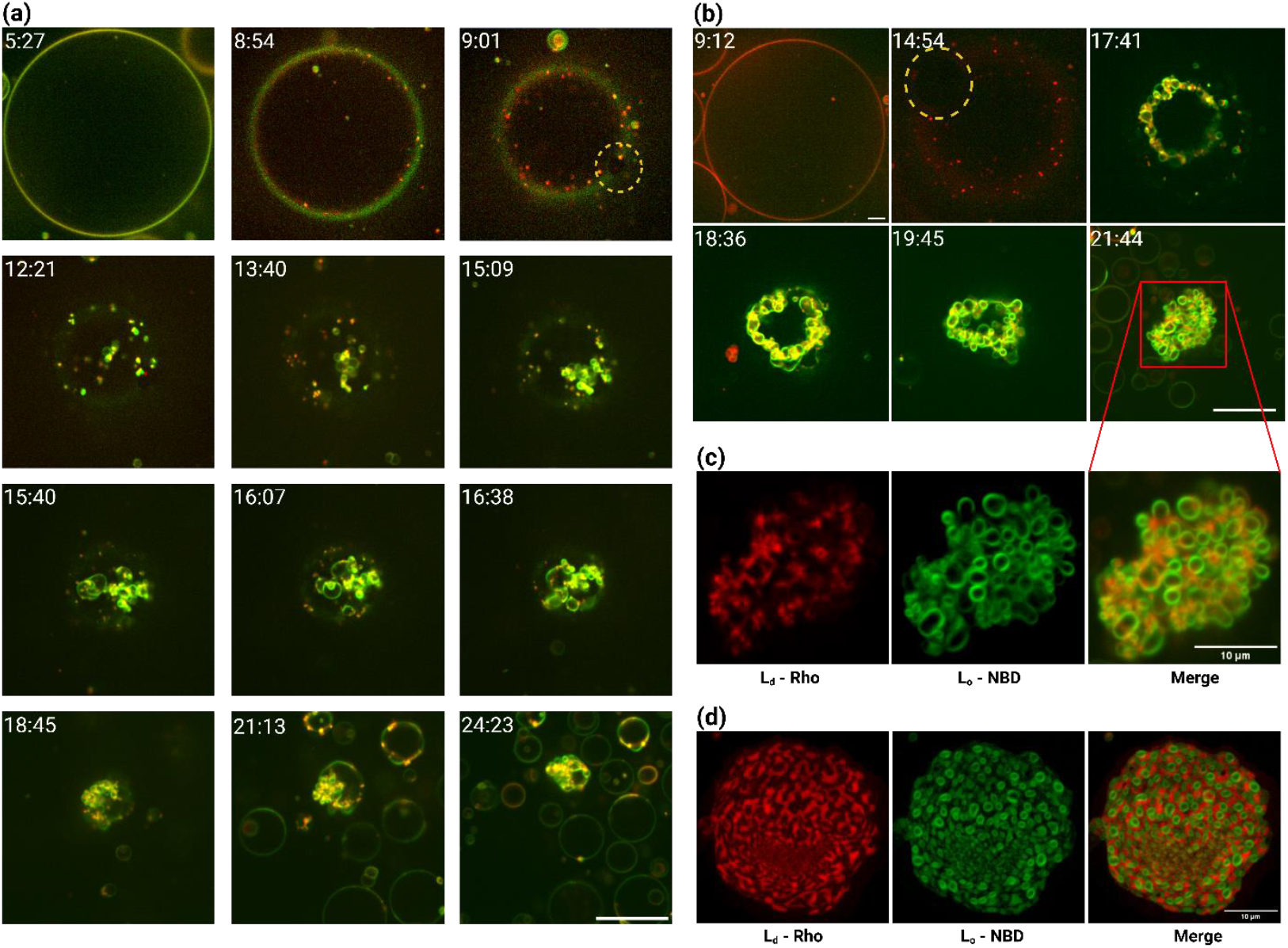
Real-time visualization of apolipoprotein interactions with GUVs containing phase-separating lipid mixtures. **A selection of confocal fluorescence microscopy images** (xy-scans) revealing vesiculation of equimolar GUVs consisting of POPC, SM and Chol and doped with 1 mol% % Rho-B-DOPE (Ld) (red) and 3 mol% % NBD-PE (Lo) (green). (a) Montage of Supplementary Video 4 depicting phase separation and vesiculation following the introduction of ∼3.5 µM Δ49ApoA-1. Scale Bar 20 µm. (b) Montage of Supplementary Video 5, showing the upper hemisphere of the vesicle. Scale bar 20 µm (**c**) Single channel and merged confocal fluorescence micrographs of a collection of vesiculated spheres from Figure 1. Clear maintained structure within the cholesterol-rich domain (green). Scale bar 10 µm.

Instances of phase separation precede large-scale morphological remodeling of the membrane. In the minutes following the appearance of phase-separated domains (12:21), structures of high-intensity NBD-PE (green) are observed to appear from the membrane. These structures appear circular and folded with an average diameter of ∼2 µm. They also seem to colocalize with the smaller Rho-DOPE domains (**Fig. 4a (13:40) & Fig. 4b (17:41)**). Over the next several minutes, formation continues, transforming the GUV into a tightly associating cluster of spherical features (24:23). These observations are fully reproducible and observed in multiple independent experiments (**Supplementary Videos 4, 5 & 6**). Across these experiments, the size of these folded structures ranges from 0.3-4.6 µm in diameter, with a density average of ∼0.2 µm^-2^ .

These localized substructures, which emerge at the membrane boundary (**Fig 4a**.), form by folding at the membrane, appear circular, and remain bound at the membrane surface – consistent with localized vesiculation. This vesiculation of multi-component GUVs reveals a striking separation of membrane lipids (**Fig. 4c**). This effect is observed most conspicuously following the vesiculation of 2:2:1 (POPC:SM:Chol) GUVs (**Fig. 4d**). Here, the NBD-PE label (green) indicates that the structured portion of the membrane domain is primarily composed of cholesterol-enriched, sphingomyelin (**Fig. 4c (L**_**o**_ **) & 4d (L**_**o**_ )), while the state of cholesterol-poor, POPC domains reveal no resolvable structure (**Fig. 4c (L**_**d**_ **) & 4d (L**_**d**_ )). Other examples of similar, ‘colony-like’ formations can be seen in **Figure S4**.

## DISCUSSION

Taken together, the results presented here reveal an array of morphological and topological transformations of giant unilamellar vesicles induced by their interactions with apolipoproteins. They show gross membrane remodeling, characterized by a well-choreographed sequence of events including the formation of transient, microscopic pores, solute leakage, and vesiculation *en route* to the reconstitution of HDL-like particles.

The formation of transient pores during the early phase of apolipoprotein – membrane interaction suggests the buildup of membrane tension while the GUV is still nominally intact (**Fig. 1a**). There are two likely scenarios that explain these changes. ***First***, it is likely that there is a loss of membrane lipids preceding optically detectable transformations of the GUV membrane. It is well established that lipid-free apolipoprotein readily recruits lipid from organized membranes (*38, 45*), and that membrane active, α-helical adopting peptides interact with phospholipid bilayers on a rapid timescale – often within a second (*49-51*).

As a consequence, NLP formation requires that lipid be drawn from the GUV membrane, reducing the lipid density, which, in turn, raises the membrane tension while the GUV is still intact (*52*). Note that this scenario only applies to the early phase when the vesicular integrity is not compromised, and that the isotonic conditions within the bath ensure the constant volume [The difference in solute concentration (𝛥C) between the bath and the encapsulated solvent is initially isotonic. Thus, the osmotic pressure acts to maintain the encapsulated volume, as prescribed by Van’t Hoff’s equation (𝛥Π = RT𝛥C), where R is the gas constant and T the absolute temperature]. Beyond a threshold lateral tension, the membrane ruptures through the well-known phenomenon of tension-induced membrane poration (*53-56*). Specifically, it is now known that when membrane tension exceeds ∼6 mN/m, energetics favor pore formation (*55*). This, in turn, sets the stage for additional morphological changes: (*57*) (i) the open pore permits the redistribution of lipids, allowing the membrane to relieve lateral tension (σ), and (ii) the encapsulated liquid (solute and solvent) leaks through the open pore due to the excess of Laplace pressure, (ΔP = 2σ/R ) decreasing the radius of the vesicle, R . These changes culminate in the healing of the pore and the resealing of the membrane. With the continued removal of lipids by ApoA-I, the process may be repeated multiple times. ***Second***, it is likely that the reversible docking of the apoproteins at the membrane surface during the early phases induces membrane tension. When the membrane is intact and moderately tense protein binding renders the membrane more tense. The ensuing poration relieves the tension and undocks the protein. Additional work is needed to resolve these mechanistic possibilities.

The observed kinetics of solute leakage are also instructive. The time taken for the encapsulated intensity of NBD-G to homogenize to the background intensity (∼20 ± 7 s) (**Fig. 1d**) is significantly longer than the lifetime of any pore formation events we observe (0.2 – 2 s). Stochastic pore nature and optical limitations of confocal microscopy limit our direct observations of transient pores to, at most, a single pore per GUV, however, we often observe secondary evidence of multiple pore formation events through fluctuations in membrane radius (**Supplementary video 2**). It seems reasonable that the leakage of the dye occurs through multiple pore formation events: multiple rounds of lipid removal and tension generation, because of the continued Δ49ApoA-I activity (**Fig. S2**). In a striking parallel, this behavior resembles that of membrane equilibration under hypotonic stress. Here, repeated cycles of elemental biophysical processes – including osmotically-mediated water influx, generation of membrane tension, membrane failure through pore formation, solute efflux through open pore, and pore healing – lead to osmotic equilibration (*54*).

The large-scale membrane morphological changes are consistent with a known aspect of ApoA-I-membrane interactions, namely conformational change. It is now well-established that the membrane binding induces a significant conformational shift in ApoA-I, increasing the helical segment from ∼50% to ∼80% (*32, 33*). The increase in the helical segment population then triggers enhanced “interfacial activity (*58*),” which reflects the increased ability of the protein to (i) bind to the membrane; (ii) partition into the membrane–water interface, acting as a wedge altering the packing and organization of the lipids within the membrane (*59, 60*); and induce an area mismatch (*61*) between the two individual leaflets that make up the phospholipid bilayer. This is further supported by the coarse-grained molecular dynamics simulations, in which ApoA-I exhibits long term association with each of the simulated membrane compositions (**Fig. 3**). Such association and dwell time would be needed to support these mechanical deformations that synergistically serve to reduce the energy cost of curvature generation (*62*) and thus promote vesiculation, as we observe.

Our coarse-grained molecular dynamics simulations (including both WT and Δ49ApoA-I) with lipid bilayers composed of 100% POPC, 4:1 POPC:POPS, 1:1:1 POPC:DPSM:Chol, and the PM8 lipid mixture (**Fig. 3 & S5**) reveal a strong composition dependence of ApoA-I interactions. The finding that the presence of negatively charged lipid species (i.e., POPS) promotes the embedding of the protein into the membrane surface is consistent with our observations of ApoE-3 and ApoE-4 interactions (**Figs 1b, S3**, and **S4**). However, the simulations also reveal only subtle morphological deformation in POPC, which is at variance with our observations of gross membrane remodeling (**Fig. 1a**). We suspect that these discrepancies arise because of the restraints placed on the double-belt model, which limit protein refolding during simulations. They further suggest that the apolipoprotein interactions with membranes composed of zwitterionic lipids alone may proceed differently and stabilize different protein conformations from those in the double belt model. But this proposition is unverified.

Further, our simulations suggest that the PM8 membrane composition induces the strongest protein-membrane interactions. By contrast, apolipoproteins introduce minimal perturbations to simpler single-component POPC membranes and even ternary mixtures containing PC, SM, and Cholesterol in our models. This then supports the alternating access mechanism (see introduction section above), in which the activated ABCA1 flips phospholipids from the cytoplasmic to the outer, exocytoplasmic leaflets of plasma membranes (*14*) and that lipids from both the leaflets participate in the HDL formation. This composition models the inner leaflet of the mammalian plasma membrane (*63-66*) – the lipids species believed to be primarily reconstituted within HDL, *in vivo* (*67, 68*).

Next, we consider the implications of the observed lipid phase separation (**Fig. 4**) during the remodeling of 1:1:1 POPC:SM:Chol GUVs upon interaction with Δ49ApoA-I. We recall that under the experimental conditions, the phase diagram predicts uniform phase with no phase-separated domains at optical length scales (*41*). There are two plausible scenarios that explain these observations. ***First***, membrane interaction of amphipathic α-helical adopting peptides have long reported generation of local membrane curvatures. Prominent examples include BAR domains (*69, 70*), GTPases (*71*), annexin B12, α-synuclein (*72*) and clathrin (*73*). Our experiments suggest that apolipoprotein interaction with a vesicular membrane leads to curvature generation (**Fig. 1a & 1b**). The physical space of the peptide acts as a wedge within membrane, supported by the presented coarse-grained MD simulations depicting ApoA-I association with the outer leaflet of varying membrane compositions (**Fig. 3a & 3b**). Insertion by the helical segments of ApoA-I into a mixed lipid membrane induces localized region of altered lipid curvature, packing and composition (*74*), reported to create microdomains capable of separating lipid species into coexisting liquid phases (*75*). A cooperative effect of ApoA-I peptides could lead to the instances of micro-scale domains, which we observe.

***Second***, it is known that HDL and ApoA-I play central roles in the removal of cholesterol from the cell membranes through reverse cholesterol transport (RCT) (*8*). Based on this consideration, it appears reasonable that the ApoA-I preferentially removes cholesterol from GUVs comprised of POPC, egg-SM, and cholesterol. It can thus tip the stoichiometric balance of the initially equimolar mixture in favor of one, which favors lipid phase separation (*76*). Either or both the cooperative effect of the α-helical peptide disruption and the selective removal of cholesterol could be responsible for the sorting of the membrane lipids into the observed membrane domains.

## ACKNOWLEDGMENTS

A.N.P & C.F.C acknowledge the preparation of WT ApoA-I, ApoE-3 and ApoE-4 for these experiments by the J.V. lab and graduate student, Maha Al-Habeeb. A.N.P & C.F.C acknowledge the microscopy facilities provided by the UC Davis LMC. T.S.C & T.N.O thank Livermore Computing and the Livermore Computational Grand Challenge for computing time. Figures were assembled, in part, utilizing the online tool Biorender.

## MATERIALS AND METHODS

### Materials

1-palmitoyl-2-oleoyl-sn-glycero-3-phosphocholine (POPC), 1,2-dioleoyl-sn-glycero-3-phosphocholine (DOPC), 1-palmitoyl-2-oleoyl-sn-glycero-3-phospho-L-serine (sodium salt) (POPS), egg-sphingomyelin, Cholesterol, and lissamine rhodamine B 1,2-dioleyl-sn-glycero-3-phosphoethanolamine (Rho-B DOPE) were acquired from Avanti Polar Lipids (Alabaster, AL). Glucose was obtained from Sigma-Aldrich (St. Louis, MO), and 2-deoxy2-[(7-nitro-2,1,3-benzoxadiazol-4-yl) amino]-D-glucose (2-NBDG) was obtained from Cayman Chemical (Ann Arbor, MI). Sucrose was obtained from EMD Chemicals (Philadelphia, PA). Chloroform, head-labeled N-(7-nitrobenz-2-oxa-1,3-diazol-4-yl)-1,2-dihexadecanoyl-sn-glycero-3phosphoethanolamine, triethylammonium salt (NBD-PE) and head labeled Texas Red™ 1,2-dihexadecanoyl-sn-glycero-3-phosphoethanolamine, Triethylammonium Salt (Texas Red™ DHPE) was obtained from Thermo Fisher Scientific (Waltham, MA). Δ49Apolipoprotein A-1 was provided by the Lawrence Livermore National Laboratory (Livermore, CA). Wildtype Apolipoproteins A-1, E-3 and E-4 were provided by the John Voss lab.

Glass-bottom, 96-well plates were obtained from Cellvis (Mountain View, CA). Indium tin oxide (ITO)-coated glass slides (5-25 Ω) were obtained from Delta Technologies, LTD (Loveland, CO). Manual Teflon-tipped syringes were obtained from Agilent Technologies (Santa Clara, CA).

### Preparation of synthetic model membranes

#### GUVs

GUV preparation via electroformation is a well-established process (*77, 78*). A lipid stock of a desired composition at a concentration of 2 mg/mL is prepared in chloroform. For single component synthetic membranes, POPC (99 mol %) is dissolved with a Rho-B DOPE label (1 mol %). Cholesterol containing membranes are prepared with a combination of POPC, egg-Sphingomyelin, and Cholesterol with the fluorescent labels of Rho-B DOPE (L_d_ domain) and NBD-PE (L_o_ domain) at a molar ratio of either 32:32:32:1:3 or 38.4:38.4:19.2:1:3, respectively. DOPC:POPS or DOPC:DOPS lipid mixtures are doped with prepared with Rho-B DOPE label at a molar ratio of 79:20:1. For all lipid stock, 15 µL is deposited onto the conductive side of two ITO-coated slides and spread evenly on half of the slide as it evaporates. These slides are further dried inside a vacuum desiccator from anywhere from 2 – 24 h. Once dry, a chamber is formed using a 1-mm-thick rubber O ring (Ace Hardware, Davis, CA) and sealed with high-vacuum grease (Dow Corning, Midland, MI). After the O ring is adhered to the conductive side of the ITO-slide, the chamber is hydrated with ∼1 mL of a 100 mM Sucrose solution. The chamber is sealed with the second, inward facing lipid cake, ITO-slide ensuring no trapped air bubbles. The two slides form the chamber in a way that the halves without dried lipid are facing in opposite directions to avoid the alligator clips from the function generator from touching both slides, effectively forming a less resistant pathway for any current. A 2.2 V(pp) AC Sine wave is applied across the two slides at a frequency of 10 Hz for 1 h, followed by another 2.2 V(pp) AC Sine wave at a frequency of 2 Hz for 30 minutes. Vesicles containing egg-SM undergo this step at a temperature of 55 °C. All samples undergo this step covered to protect from light. After GUV formation, the chamber is disassembled, and the solution is pipetted into a small centrifuge tube and stored at 4 °C. Vesicles prepared are either used or discarded within a week of preparation.

### Preparation of Δ49ApoA-I and WT Apolipoprotein

#### Cell-free synthesis of Δ49ApoA-I protein

Cell-free reactions were set up using RTS 500 ProteoMaster *E. coli* HY Kit (Biotechrabbit GmbH, Hannover, Germany). Reaction components (lysate, reaction mix, feeding mix, amino acid mix, and methionine) were combined as specified by the manufacturer. Each 1 mL reaction contained 15 µg Δ49ApoA-I plasmid DNA and 20 µL Fluoro-Tect™ GreenLys tRNA (Promega, Madison, USA). The reactions were incubated at 30 °C, with shaking at 300 rpm for 18 h in a floor shaker. After the reaction, ApoA-I protein is purified using nickel affinity chromatography. Briefly, 1 mL of 50% slurry complete His-Tag Purification Resin (Roche Molecular Diagnostics, Basel, Switzerland) was equilibrated with equilibration buffer (50 mM NaH_2_ PO_4_, 300 mM NaCl, pH 8.0) with 10 mM imidazole (Sigma-Aldrich, St Louis, MO) in a 10 mL chromatography column. The total cell free reaction was mixed with the equilibrated resin and was incubated/nutated at 4 °C for 1 h. The column was then washed 6 times with 1 mL wash buffer containing (50 mM NaH_2_ PO_4_, 300 mM NaCl, 20 mM imidazole, pH 8.0). ApoA-I is eluted with 6 x 300 µl elution buffer (50 mM NaH_2_ PO_4_, 300 mM NaCl, 250 mM imidazole, pH 8.0). All elutions were analyzed by SDS-PAGE and peak fractions containing labeled protein were combined. Pooled fractions were dialyzed in PBS (pH 7.4) and then stored at 4 °C. Protein levels were quantified using a Qubit protein test according to manufacturer’s instruction (Thermo Fisher Scientific, Carlsbad, CA).

### Production and Purification of ApoA-I protein

Human ApoA-I was expressed in *Escherichia coli* strain BL21 Star (DE3)pLysS cells (Life Technologies, Carlsbad, CA, USA) from the human ApoA-I gene containing a hexa-His affinity tag at the N-terminus. The ApoA-I gene was cloned into the pEXP-5 plasmid (Novagen, Inc, Madison, WI, USA) and transferred into the bacteria and cultivated at 37° C in LB medium with 50 µg/mL of ampicillin and 34 µg/mL of chloramphenicol. Protein expression was induced for 3-4 h following the addition of 0.5 mM isopropyl-beta-thiogalactopyranoside (Sigma, St Louis, MO, USA). Following cell disruption, ApoA-I was purified from the soluble fraction of the cells using a His-Trap-Nickel-chelating column (GE Healthcare, Uppsala, Sweden) and a mobile phase of phosphate-buffered saline (PBS), pH 7.4 with 3 M guanidine. The protein was washed in PBS (pH 7.4) containing 40 mM imidazole and then eluted with PBS containing 500 mM imidazole. Imidazole was removed from the protein sample by dialysis (10K MWCO Slide-A-Lyzer, ThermoFisher) equilibrated with PBS, pH 7.4. Protein purity was analyzed by SDS-PAGE and concentration was determined using a Nanodrop 2000c spectrophotometer (Thermo scientific, Waltham, MA, USA), which was based on the extension coefficient (32,490 l mol^-1^ cm^-1^ ) and molecular weight (28,279 Da) of ApoA-I.

### Production and purification of ApoE protein

Human ApoE-3 or ApoE-4 protein was expressed in *Escherichia coli* cells and purified as described previously (*79*). Briefly, one liter of cells in LB broth were grown to mid-log phase at 37 °C. These were then induced with 0.1% arabinose, and incubated for an additional 4h at 37°C. The cells were harvested by centrifugation, and inclusions bodies isolated and washed as described previously (*80*). To purify the protein, washed inclusion bodies were dissolved in 8 M urea, 200 mM NaCl, and the sample filtered through a 0.2 µm filter and then chromatographed on a SuperDex 200 (Pharmacia) size exclusion column using a mobile phase of 8M urea, 200 mM NaCl, 10mM TRIS (pH 7.5) and 1mM EDTA. The fraction was then desalted using a SuperDex 200 column using a mobile phase of 8M urea, 10mM TRIS (pH 7.5) and 1mM EDTA and then bound to a mono-Q ion exchange column. The column was washed with 8M urea,10 mM TRIS (pH 7.5) and 1mM EDTA, and the protein eluted using a gradient of NaCl. The protein was then refolded by dialysis (Slide-A-Lyzer, ThermoFisher) against 0.1M ammonium bicarbonate and then PBS buffer pH 7.4. The sample was concentrated using centrifugal spin concentrators with a molecular size cutoff of 30 kDa (Millipore), and the protein concentration determined using the Pierce BCA kit (ThermoFisher, USA).

### ApoA-I incubation assays

#### GUV incubation with Apoprotein

All spinning-disk confocal fluorescence imaging was taken in conjunction with glass-bottom, 96-well plates (Cellvis). Experiments observed in this way have a final volume of 100 µL per well. 2 µL of GUV solution prepared in a 100 mM sucrose solution are added to a bath of osmotically balanced, 100 mM Glucose solution. The difference in density of the GUVs allows them to settle on the bottom of the well. Control experiments are imaged in these conditions, otherwise 1-5 µM Δ49ApoA-I, WT ApoA-I or ApoE-4 is added by micropipette slowly (3-5 s) and gently to the center of the well.

### GUV leakage assay

GUVs containing soluble dye are prepared as described above; Slides are hydrated with a 100 mM Sucrose & 100 µM of 2-NBDG dye before the electroformation step. Both POPC and cholesterol containing GUVs are prepared in this way. When observing the leakage of dye from the inside of GUVs, incubation with ApoA-I follows the above-described method. When observing the leakage of dye from extracellular space, the bath is instead prepared with 100 mM Glucose & 100 µM 2-NBDG followed by the introduction of GUVs prepared in only 100 mM Sucrose. Spinning-disk confocal fluorescence imaging with 488 nm & 561 nm lasers were used simultaneously to observe any ApoA-I-induced leakage between the intra- and extracellular space. The difference in fluorescence intensities between these spaces is used for quantification of the leakage over time.

### Coarse-grained molecular dynamics simulations

The sequence of human ApoA-I was taken from Uniprot database (Entry P02647 (*81*)) and used as an input to *Nanodisc Builder* (*82*) in order to generate an atomistic structural model for the ApoA-I double-belt, referred as WT. The mutant sequence where first 49 residues were deleted was also fed into *Nanodisc Builder* to generate a model for the Δ49ApoA-I double-belt, referred as Δ49. Next, the atomistic models were converted to coarsegrained structural models using *martinize2* with Martini 2.2 force field parameters (*83, 84*). The elastic network was added to the structures with a force constant of 900 kJ/mol/nm^2^ and a lower (upper) distance cutoff of 0.5 nm (1.2 nm). A 30 nm x 30 nm POPC, 4:1 POPC:POPS, 1:1:1 POPC:DPSM:CHOL, or PM8 bilayer was built with *Insane* (*85*) and solvated with 0.15 M NaCl and 10% antifreeze water (WF) beads; these systems included the coarsegrained WT and Δ49 belt proteins placed 4 nm away from the membrane. PM8 membrane was set as a symmetric bilayer with the lipid composition of the inner leaflet composition of a typical mammalian plasma membrane model used in a previous study (*86*) and included 13.9% POPC, 7.5% PAPC, 5.4%POPE, 16.1% DIPE, 10.8% DPSM 28% CHOL, 16.1% PAPS and 2.2% PAP6.

For each membrane composition, we carried out three independent simulations for 10 µs, 5 µs and 5 µs using the protocol presented by Ozturk et al. (*87*). We carried out these simulations twice: the one set with restraints on the ApoA-I belt dimers and the other set without any restraints. The first set had two distance restraints that were only in effect if the distances became smaller than 8 nm in WT ApoA-I and 6 nm Δ49, respectively, with a force constant of 1000 kJ/mol/nm^2^ . Two distance variables were defined using the geometric centers of the residues (1) 110-112 and 250-252 and (2) 280-282 & 410-412 for the WT ApoA-I model and (1) 80-82 and 190-192 and (2) 300-302 and 410-412 for the Δ49 ApoA-I model. These restraints did not allow for full closing-up of the belt but still allowed for large conformational changes. The temperature and pressure were kept at 310 K and 1 bar with v-rescale thermostat (*88*) and c-rescale barostat (*89*) during the production runs as indicated in a previous work (*87*) and are listed in the Supplementary Table 1. Note that LINCS order and iteration were set to 12 and 2 which are crucial when cholesterol is modeled with Martini 2.2 force field parameters with a time step of 20 fs (*90*). As suggested in a recent study, to avoid unphysical distortions in the membrane, the outer cutoff and timesteps between neighbor list updates were set to 1.35 nm and 20 (*91*). All coarse-grained molecular dynamics simulations were carried out with Gromacs 2024.4 (*92*) and the restraints were applied using the Colvars module (*93*). The simulation snapshots were generated with VMD (*94*).

### Characterization

#### Fluorescence microscopy

Spinning-disk confocal imaging and measurements were collected on an Intelligent Imaging Innovations Marianas Digital Microscopy Workstation (3i, Denver, CO) with attached CSU-X1 spinning-disk head (Yokogawa; Mushaninoshi, Tokyo, Japan), a Quantem512SC EMCCD camera (Photometrics, Tucson, AZ) and a Flash 4 ORCA-sCMOS camera (Hamamatsu, Hamamatsu-city, Japan). A Zeiss Plan-Fluor 63x (NA 1.4), oil immersion objective (Carl Zeiss, Oberkochen, Germany) was used in the collection of these data. Rho-B DOPE (Ex/Em 560:583) was exposed with a 50 mW 561 nM laser line, while NBD-PE (Ex/Em 463:536) and 2-NBDG (Ex/Em 465:540) were exposed with a 50 mW 488 nM laser line. Image processing was done using Fiji a public-domain software; https://imagej.net/software/fiji/), Slidebook digital microscopy imaging software (3i), and MatLab (The Math Works, Inc. MATLAB Version 2023a).

### Size Exclusion Column Purification (SEC)

The NLPs are analyzed and purified via size exclusion chromatography (SEC) with a semi-preparative Superose 6 10/300 column (GE Healthcare). SEC was run on a Shimadzu LC-20 HPLC system (Shimadzu) at flow rate 1mL/min with PBS as running buffer. NLP peak was assessed using absorption at 280 nm and analyzed with instrument software LC Solutions (Shimadzu).

### High-Speed AFM characterization of reactant NLPs

A high-speed atomic force microscope (RIBM, Japan) was equipped with a small cantilever (Ultra-Short Cantilevers (USC, NanoWorld): spring constant, k = 0.15 N/m, resonance frequency, f = 1200 kHz in water) and was operated in tapping mode at room temperature. The free oscillation amplitude was 0.9 ∼ 1.3 nm, and the typical setpoint amplitude was 85% of the free oscillation amplitude. For the sample stage, a freshly cleaved mica disk with a diameter of 1.5 mm was fixed on a glass rod with a diameter of 1.5 mm and a height of 2 mm using epoxy glue. The SEC-purified NLPs was diluted 20x in PBS buffer, and 2ul of the diluted solution was deposited onto the freshly cleaved mica and incubated for 10 minutes. The scanner was then mounted above the sample chamber with the cantilever immersed in PBS buffer. HS-AFM images were processed using ImageJ (National Institutes of Health, Bethesda, MD, USA). The height and width dimensions of each nanodisc were analyzed with Gwyddion software (*95*) and the data was plotted with Origin.

