## Supplementary Figures for "Apolipoprotein interaction induces shape remodeling and lipid phase separation in giant unilamellar vesicles"

### *Supplementary Table 1: Molecular dynamics simulations*

| **Membrane Composition** | **WT APOA1** | | **D49 APOA1** | |
|---|---|---|---|---|
| | *Without restraints* | *With restraints* | *Without restraints* | *With restraints* |
| POPC | 10 µs | 5 µs | 10 µs | 5 µs |
| POPC | 5 µs | 5 µs | 5 µs | 5 µs |
| POPC | 5 µs | 5 µs | 5 µs | 5 µs |
| POPC:POPS | 10 µs | 5 µs | 10 µs | 5 µs |
| POPC:POPS | 5 µs | 5 µs | 5 µs | 5 µs |
| POPC:POPS | 5 µs | 5 µs | 5 µs | 5 µs |
| CHOL:DPSM:POPC | 10 µs | 5 µs | 10 µs | 5 µs |
| CHOL:DPSM:POPC | 5 µs | 5 µs | 5 µs | 5 µs |
| CHOL:DPSM:POPC | 5 µs | 5 µs | 5 µs | 5 µs |
| PM8 | 10 µs | 5 µs | 10 µs | 5 µs |
| PM8 | 5 µs | 5 µs | 5 µs | 5 µs |
| PM8 | 5 µs | 5 µs | 5 µs | 5 µs |

*Supplementary Table 1: List of 48 coarse-grained MD simulations carried out in this work*

### *Supplementary Figure 1: Pore formation events*

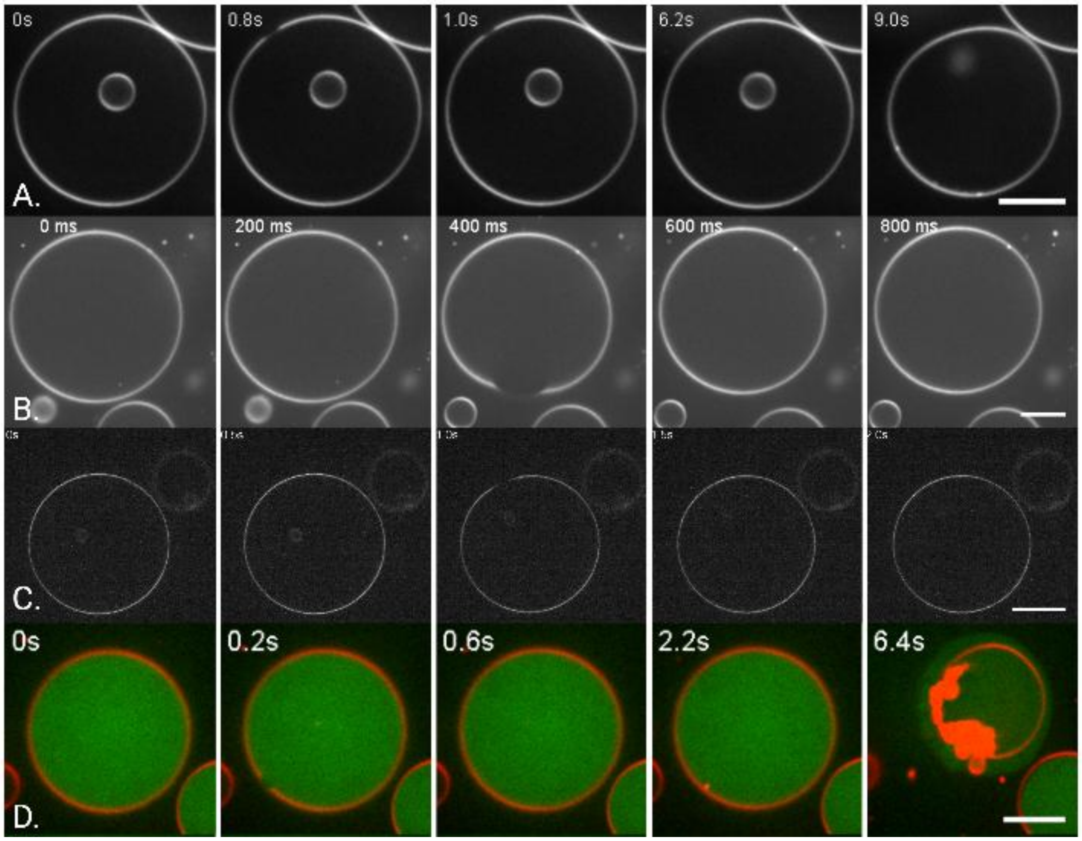


*Supplementary Figure 1: Montage of transient pore events in GUV, along with the radius of the pore ($R_p$) and the lifetime of the pore ($L_p$).* ***(a)*** *POPC GUV introduced to 0.88µM of Δ49ApoA-1. $R_p$ = ~1µm and $L_p$ = ~1.2s.* ***(b)*** *POPC GUV introduced to 0.88µM of Δ49ApoA-1. $R_p$ = 6µm and $L_p$ = ~0.2s.* ***(c)*** *1:1:1 RAFT GUV introduced to 0.44µM of Δ49ApoA-1. $R_p$ = 3.7µm and $L_p$ = ~0.5s.* ***(d)*** *POPC GUV introduced to 0.44µM of Δ49ApoA-1. $R_p$ = 1.1µm and $L_p$ = ~2s. Scale bars are 10 µm.*

#### *Supplementary Figure 2: Solute leakage through a cascade of pore formation events*

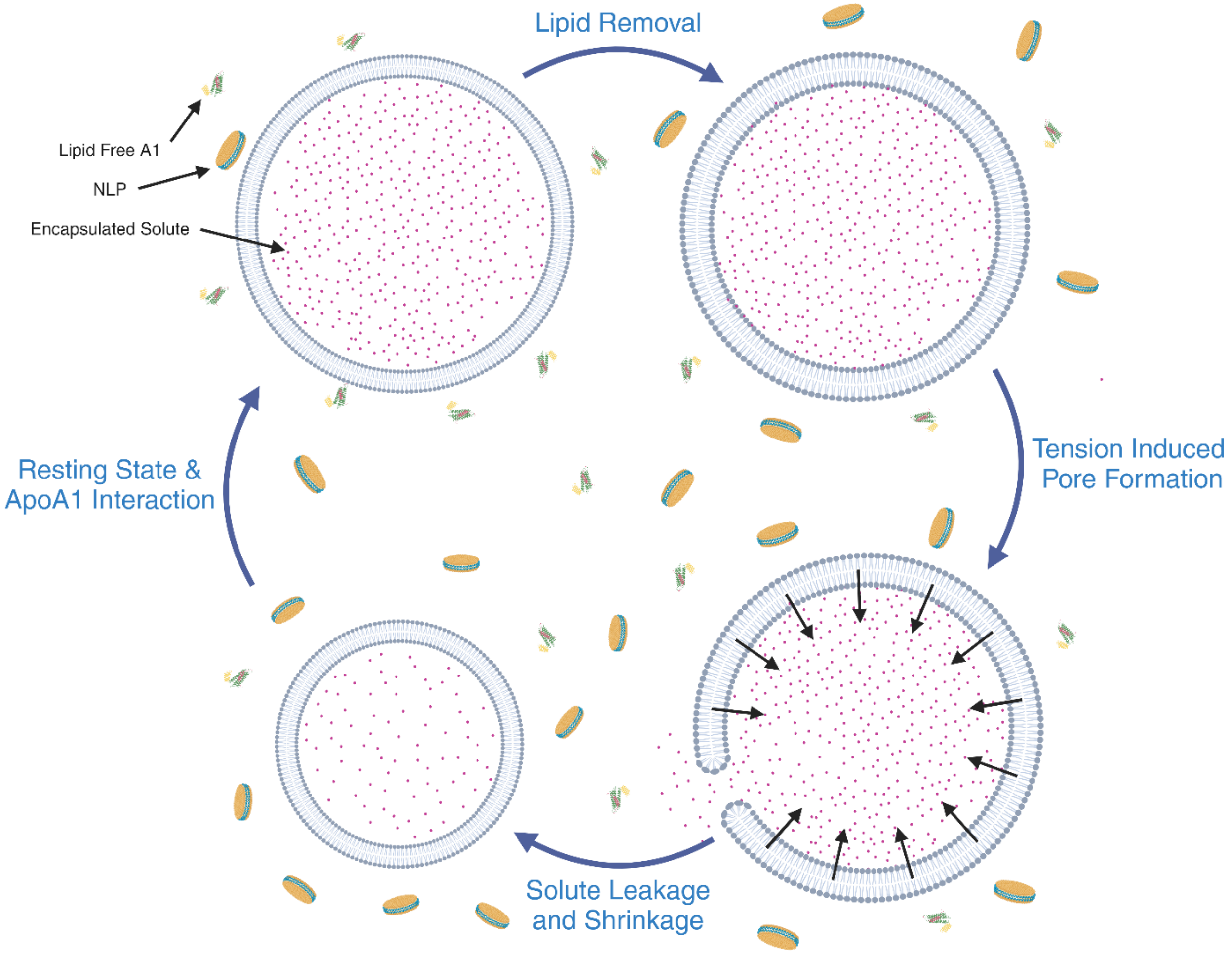


*Supplementary Figure 2: Diagram of cyclic behavior of giant unilamellar vesicles (GUVs) in response to apolipoprotein interaction. Starting from the top left, (i) an initially isotonic GUV is introduced to lipid-free apolipoprotein, (ii) upon membrane interaction, NLP formation ensues through the stochastic removal of lipid from the membrane. (iii) The GUV now experiences a reduced lipid density (an initially isotonic condition acts to maintain encapsulated volume, as solute is membrane impermeable) and increasing membrane tension until a critical threshold is reached and a transient pore is formed. This pore allows for the exchange of solute and solvent, while excess Laplace pressure assists in the resealing of the pore around a smaller volume. (iv) The GUV is now at a new equilibrium, in which continued lipid removal may ensue, and repeat this cycle.*

### ***Supplementary Figure 3: DOPC:POPS (4:1) Vesiculation due to WT ApoE-4***

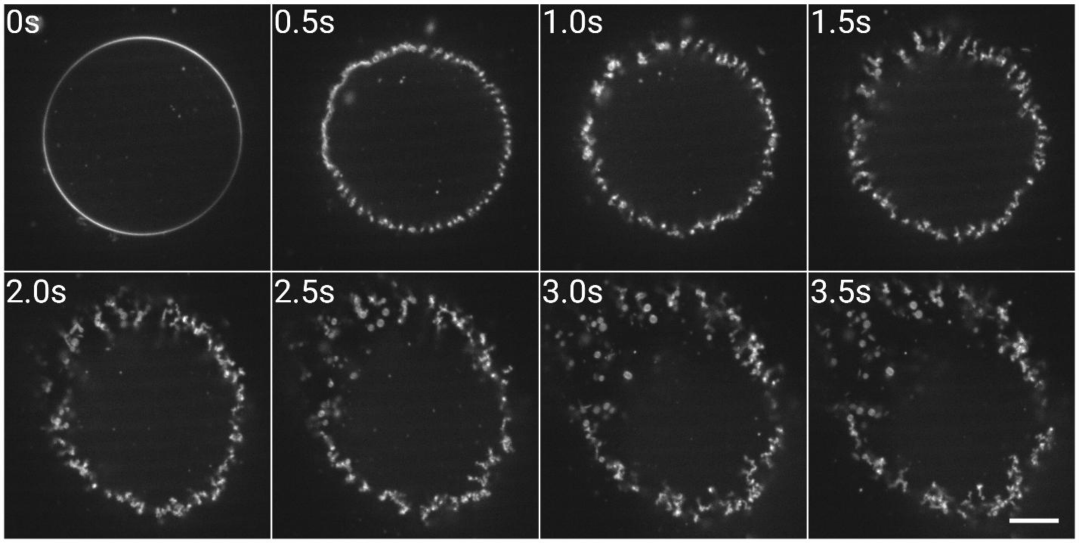


*Supplementary Figure 3: Vesiculation montage of DOPC:POPS (4:1) GUV exposed to 3µL of 0.5mg/mL WT ApoE4. Scale bar = 10 µm.*

### *Supplementary Figure 4: DOPC:DOPS (4:1) Vesiculation due to ApoE-3*

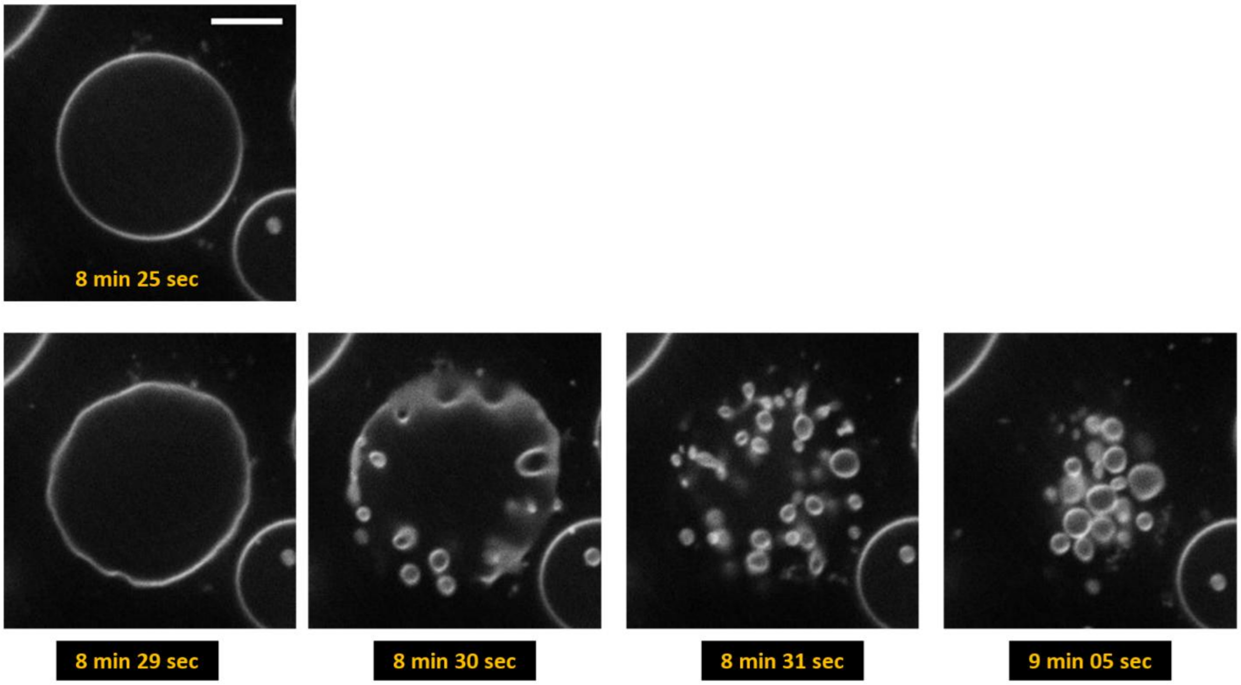


*Supplementary Figure 4: Montage of DOPC:DOPS (4:1) GUVs incubated in 7.38 μM ApoE3. Scale Bar is 10um*

### *Supplementary Figure 5: Coarse-Grained MD Snapshot of Δ49 & WT ApoA-I*

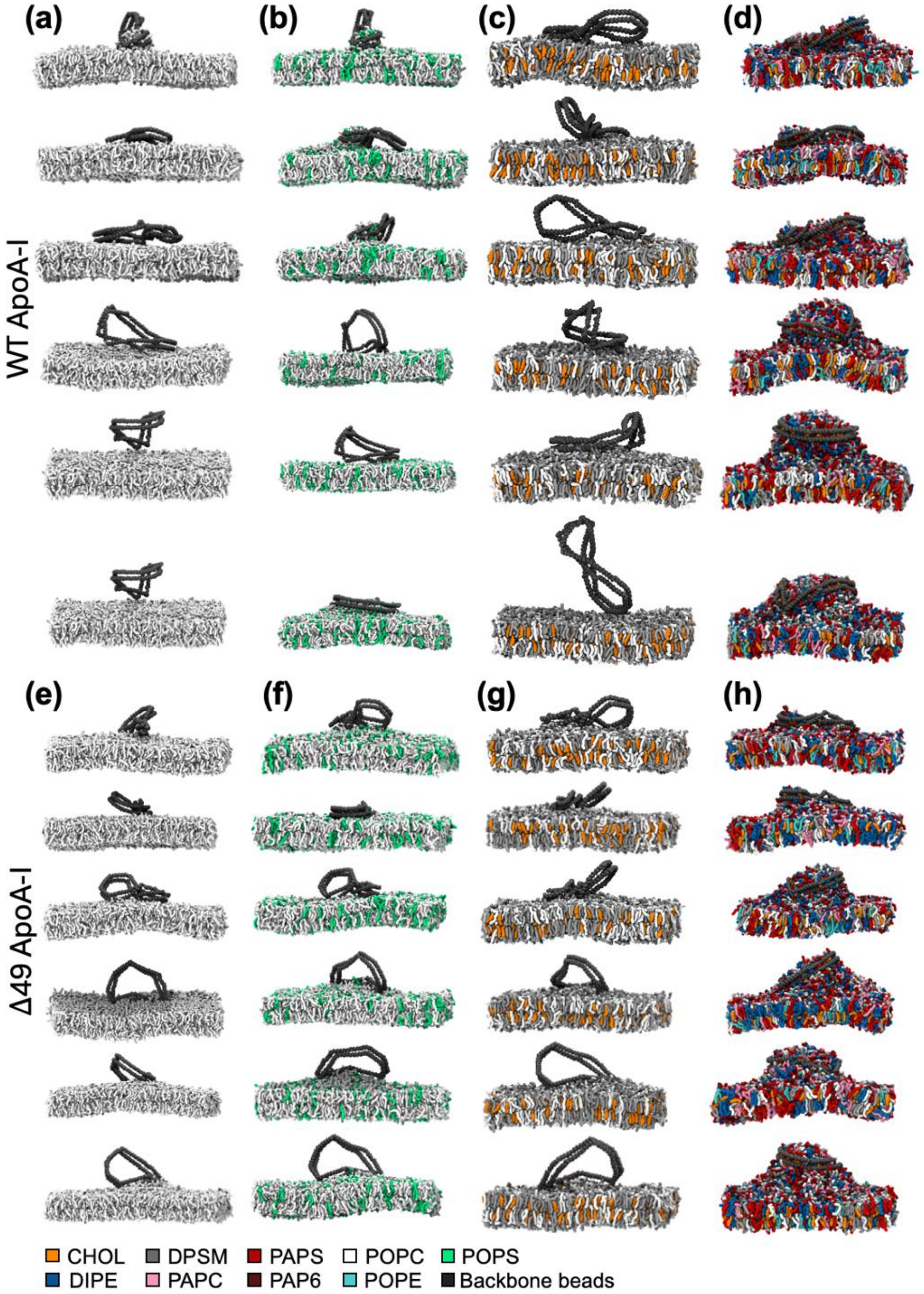


***Figure 5:*** *Last snapshots of all coarse-grained molecular dynamics simulations carried out using a pre-assembled WT or Δ49 ApoA-I double belt protein with different membranes. Panels a-d show the WT ApoA-I double-belt and the* ***(a)*** *POPC,* ***(b)*** *4:1 POPC:POPS,* ***(c)*** *1:1:1 CHOL:DPSM:POPC and* ***(d)*** *PM8 (plasma membrane mimetic model) lipid bilayers at the end of six independent coarse-grained molecular dynamics simulation. Panels e-g represent the Δ49 ApoA-I protein and the* ***(e)*** *POPC,* ***(f)*** *4:1 POPC:POPS,* ***(g)*** *1:1:1 CHOL:DPSM:POPC and* ***(h)*** *PM8 membranes. In each panel, the top row shows the simulated system at the end of 10 μs whereas the rest of the rows show the same system at the end of 5 μs-long independent simulations. The top three rows in each system show simulations are carried out without any restraints in the protein while the last three rows show systems simulated with restraints in the protein as explained in the Methods. Lipids are color-coded: CHOL in orange, DIPE in blue, DPSM in gray, PAPC in pink, PAPS in red, PAP6 in dark red, POPC in white, POPE in cyan and POPS in green. Water and ions are not shown for clarity. Backbone beads of the protein are colored in dark gray.*

### *Supplementary Figure 6: Post-vesiculation of 2:2:1 (POPC:SM:Chol) GUVs*

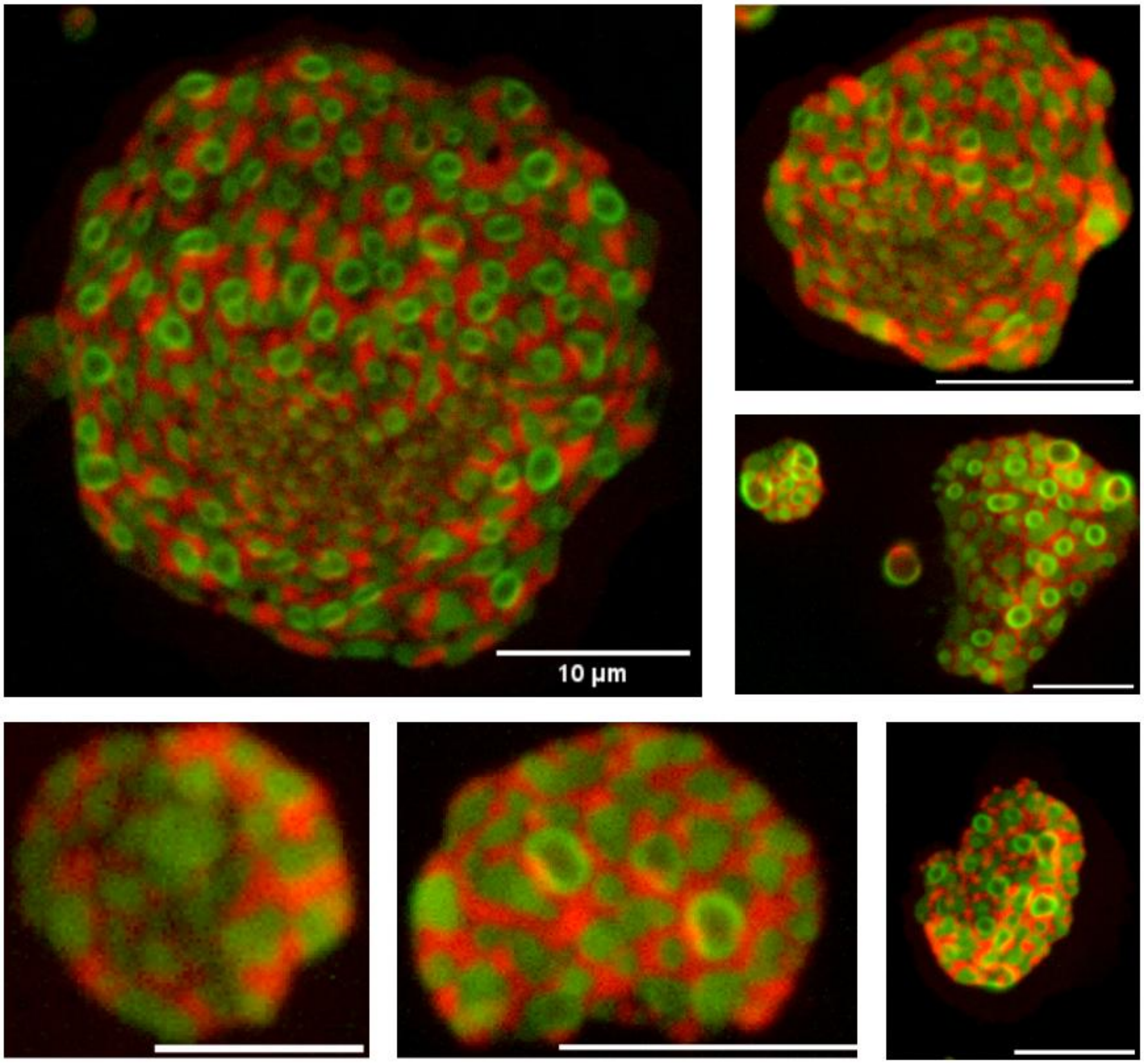


*Supplementary Figure 6: Post Vesiculation of 2:2:1 (POPC:SM:Chol) GUVs following the introduction of ~1.7µM of Δ49ApoA-1. These GUVs are doped with 1 mol % of Rho-DOPE (red) and 3 mol % NBD-PE (green). Phase separated GUVs burst rapidly upon interaction with Δ49ApoA-1. Adhered to the glass substrate, a clear distinction between membrane domains is observed. The Chol-rich, SM domain (green) retains spherical structure.*

### *Supplementary Video 1: POPC Leakage and Vesiculation*

*Supplementary Video 1: A collection of POPC GUVs doped with 1 mol % Rho-B-DOPE (red) and encapsulating 100µM of a soluble, membrane impermeable dye NBD-G (green). Upon introducing 0.88uM of Δ49ApoA-1, GUV leakage, transient pore formation and vesiculation are observed.*

### *Supplementary Video 2: POPC Leakage and Vesiculation Extended*

*Supplementary Video 2: A collection of POPC GUVs doped with 1 mol % Rho-B-DOPE (red) and encapsulating 100µM of a soluble, membrane impermeable dye NBD-G (green). Upon introducing 0.88uM of Δ49ApoA-1, GUV leakage, transient pore formation and vesiculation are observed. These effects propagate throughout the sample over the observed time (17 min).*

### *Supplementary Video 3: DOPC:DOPS (4:1) GUV Vesiculation*

*Supplementary Video 3: DOPC:POPS (4:1) GUVs doped with 1 mol % Rho-B-DOPE (red). Tubules within the GUV are a common artifact of the electroformation technique and are unrelated protein interaction. Upon introducing 3uL of 0.078mg/mL of WT ApoA-I, vesiculation of the GUV occurs.*

### *Supplementary Video 4: 1:1:1 (POPC:SM:Chol) GUV Phase separation & Vesiculation Ex. 1*

*Supplementary Video 4: Equimolar GUVs composed of POPC, egg-SM and cholesterol is doped with 1 mol % Rho-B-DOPE ($L_d$) (**red**) and 3 mol % NBD-PE ($L_o$) (**green**). Phase separation and vesiculation following the introduction of ~3.5 µM Δ49ApoA-I. At 8:40, brightness/contrast correction is applied so membrane deformation is clearly visible. A pore can be seen to appear beginning at 9:01. Scale Bar 20µm. Scale bar 10µm.*

### *Supplementary Video 5: 1:1:1 (POPC:SM:Chol) GUV Phase separation & Vesiculation Ex. 2*

*Supplementary Video 5: Equimolar GUVs composed of POPC, egg-SM and cholesterol is doped with 1 mol % Rho-B-DOPE ($L_d$) (**red**) and 3 mol % NBD-PE ($L_o$) (**green**). Phase separation and vesiculation following the introduction of ~1.76 µM Δ49ApoA-I.*

### *Supplementary Video 6: 1:1:1 (POPC:SM:Chol) GUV Phase separation & Vesiculation Ex. 3*

*Supplementary Video 6: Equimolar GUVs composed of POPC, egg-SM and cholesterol is doped with 1 mol % Rho-B-DOPE ($L_d$) (**red**) and 3 mol % NBD-PE ($L_o$) (**green**). Phase separation and vesiculation following the introduction of ~1.76 µM Δ49ApoA-I. Scale bar is 10µm.*
